# Molecular basis for regulation of human potassium chloride cotransporters

**DOI:** 10.1101/2020.02.22.960815

**Authors:** Ximin Chi, Xiaorong Li, Yun Chen, Yuanyuan Zhang, Qiang Su, Qiang Zhou

**Author notes:** These authors contributed equally to this work (Ximin Chi, Xiaorong Li, Yun Chen, Yuanyuan Zhang).

## Abstract

The Solute Carrier Family 12 (SLC12) encodes electroneutral cation-chloride cotransporters (CCCs) that are fundamental in cell volume regulation and chloride homeostasis. Dysfunction of CCCs engenders abnormality in renal function and neuro-system development. Here we presented structure of the full length human potassium-chloride co-transporter 2 (KCC2) and 3 (KCC3), the KCC3 mutants in phosphorylation mimic (P-mimic) and dephosphorylation mimic (DP-mimic) status, and KCC3 in complex with [(DihydroIndenyl)Oxy] Alkanoic acid (DIOA), a specific inhibitor of KCCs, at resolution of 2.7 Å - 3.6 Å. A small N-terminal loop is bound at the intracellular vestibule of the transport path, arresting the transporter in an auto-inhibition state. The C-terminal domain (CTD) of KCCs is structure-solved for the first time, revealing two conserved phosphorylation harboring segments (PHSs), which exhibit different conformation between P-mimic and DP-mimic mutants, explaining the inhibitory effect of phosphorylation. DIOA is located in between the two transmembrane domains, tightly bound to the loop between TM10 and TM11, locking the transporter in inward-facing conformation. Together, our study makes important steps in understanding the sophisticated regulation mechanisms of KCCs, with prospect for the specific drug development.

## Introduction

CCCs mediate the electroneutral across cell membrane transport of sodium and/or potassium cations and chloride anion. Two major classes of CCCs exist in mammals, the sodium-independent co-transporters consisting of KCC1-4 and the sodium-dependent co-transporters including NKCC1-2 and NCC (1–3). Phosphorylation regulates the transporter activity along with a series of kinases, e.g. WNK-SPARK/OSR1 system (4, 5). The dephosphorylation and phosphorylation mutations activate and inactive KCCs, respectively (6–9), though activated phosphorylation site was also identified in KCC2 (10). Under the hypertonic environments, the phosphorylated NKCCs are activated to facilitate the ion influx into the cell, while the phosphorylated KCCs are inhibited. When cell is faced to the hypotonic circumstances, dephosphorylated KCCs induce the potassium and chloride ions efflux to the extracellular space, promoting cell shrink to the normal volume (11). Disorder of the SLC12 family results in hereditary diseases including epilepsy (12, 13), Anderman syndrome (14), Gitelman syndrome (15), Bartter syndrome (16), and deafness (17, 18).

KCC2, which was identified in 1996 from rat brain cDNA library, is restrictedly expressed in central nervous system (19). Distinctive from other KCCs, KCC2 is reported to be isotonicity active (20). The extrusion of chloride ion mediated by KCC2 provides the basis for GABA-induced neuron inhibition (21). Inhibitory phosphorylation of T906 and T1007 in KCC2 is reported to be fundamental in the nervous system development (7–9). KCC3 was identified in 1999 by three groups (22–24), which is expressed in multiple tissues and organs including brain, kidney, and muscle. The CTD of KCC3 is reported in the interaction with brain-specific creatine kinase; deficiency of the interaction is correlated to hampered KCC3 activity and agenesis of corpus callosum (25). NTD of KCC3 also harbors functional key points (26, 27). The same inhibitory phosphorylation sites are conserved in KCC3, ablation of phosphorylation lead to co-transporter activation and neuron abnormality (6, 28).

KCC2 and KCC3 are important therapeutic targets in the case of peripheral neuropathy (29), epilepsy (30), and acute central nervous injury (31). The widely-applied diuretics furosemide and bumetanide favors NKCCs than KCCs (32). Although some KCCs specific inhibitors were identified (33–36), there is still a lack of specific drugs for KCCs (37). Although most diuretics are hepatotoxic in clinical treatment, drugs with poor specificity can still produce side-effects considering the wide distribution and overall structural similarity between CCCs (38).

The recently reported *Dr*NKCC1 and KCC1 structures provide the structural glimpse for the CCCs family (39, 40). However, the limited sequence similarity between *Dr*NKCC1 and KCCs as well as the lack of the cytosolic domain in the reported KCC1 structures hinder the elaboration of the KCCs working mechanism in detail. Here, we present single particle cryo electron microscopy (cryo-EM) structures of the full-length human KCC2 and KCC3 at resolution of up to 2.9 Å. The structures are very similar to each other, existing as a domain-swapped dimer. The CTD of the KCC transporters rotates around 70° compared with *Dr*NKCC1, revealing structural difference among CCC family. An auto-inhibitory N-terminal loop was observed located in the entry of the cytosolic vestibule and function as a blocker. The PHS is a flexible segment located in the peripheral region of the CTD, which adopts different conformations compared with their counterpart in *Dr*NKCC1. In order to investigate the effect of phosphorylation, P-mimic and DP-mimic mutants of KCC3 were generated and structure-solved. Structural analysis suggests that phosphorylation inhibits the transport activity by stabilizing the inward-facing conformation through PHSs. DIOA, a selective inhibitor of KCCs (36), is bound between the transmembrane domains of the two protomers, exerting it inhibitory efficacy also by stabilizing the inward-facing conformation of the transporter. Our studies provided structural basis for the regulation mechanisms of KCCs by both *cis*-elements including N-terminal loop and PHSs, and *trans*-element, DIOA, shedding light for the development of specific drugs for KCCs.

## Results

### Purification and structure determination of human KCC2 and KCC3

In order to understand the regulation mechanism of KCCs, we over-expressed the full-length human KCC2 and KCC3 in HEK293F cells and purified the proteins. Then the protein samples were concentrated and subject to structural determination by single particle cryo-EM. Finally, we solved the structures of KCC3 and KCC2 at overall resolution ranging from 3.3 Å to 3.2 Å. The local resolution of the TM/extracellular domains (ECD) and CTD can be further improved to 3.1 Å and 2.9 Å for KCC3 or to 2.9 Å for KCC2 by focused refinement, respectively, which allows the unbiased model building (Figure 1A, 1B, Supplemental Figure S1, S2, Table S1). Details of protein purification, cryo EM sample preparation, data collection, image processing and model building can be accessed in the Supplemental Information.

**Figure 1.**
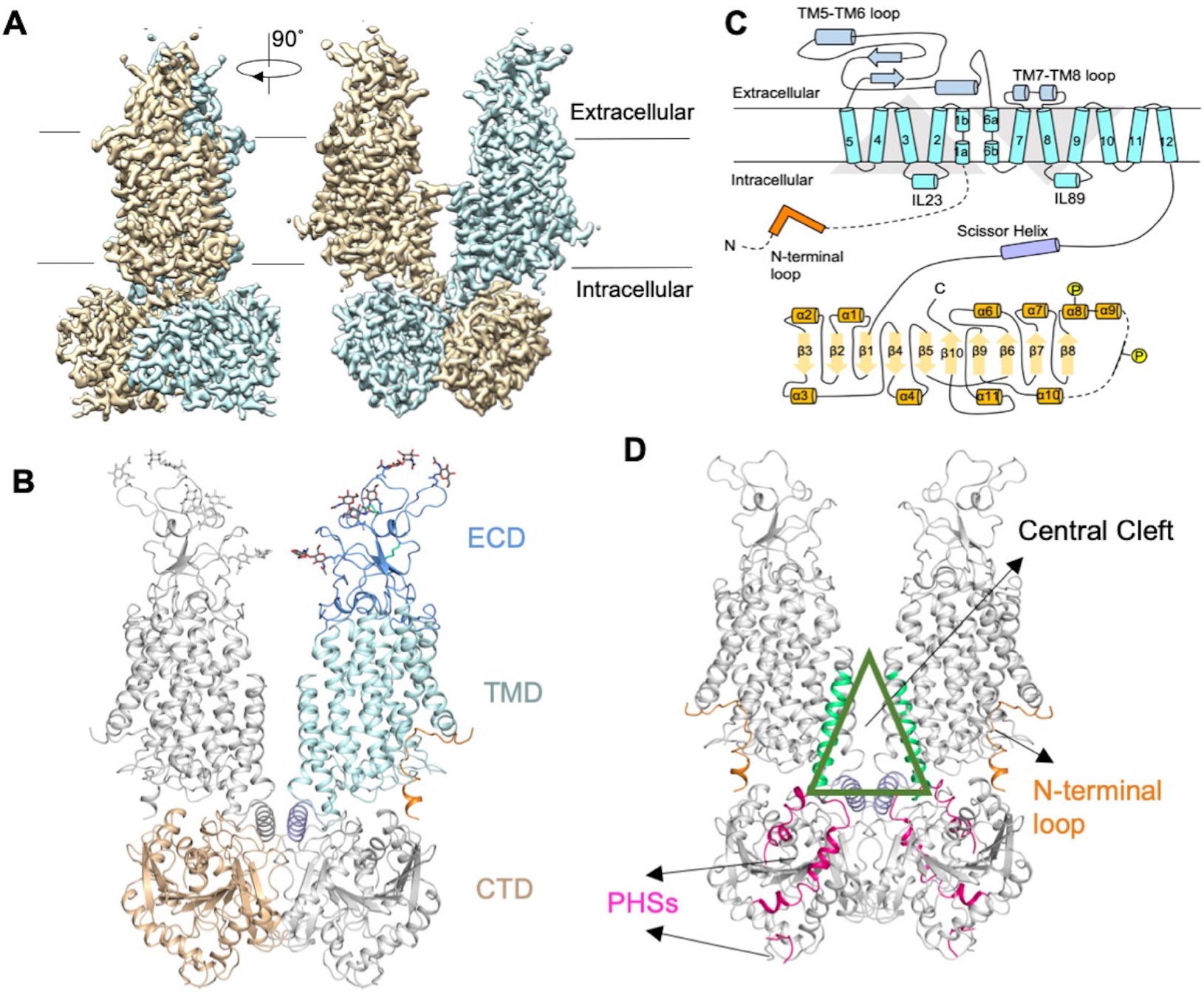
Overall structure of human KCC3. (A) Cryo-EM map of full length human KCC3, generated by merging maps focused refined on TM/ECD and CTD. (B) The structure model of human KCC3 colored by domains. Carbohydrate chains were shown in sticks. Unless otherwise indicated, the same domain color scheme is applied to all figures. (C) Topology diagram of human KCC3. only one protomer is shown. (D) Summary of the key regulatory elements or region discussed in this study. Center Cleft is formed by TM12 and Scissor Helixes. N-terminal loops are located in the cytosolic entry of TM domains. PHSs are located in CTD.

The structures of KCC2 and KCC3 are very similar to each other with root mean squared deviation of 1.39 Å over 841 Cα atoms. In next text, we mainly focused on KCC3 for structural analysis. The solved KCC3 structure covers ~85% of the sequence of KCC3. Based on the cryo-EM map and mass spectra results, phosphorylated threonine was built on T940 and T997. The region of Q945-R960 is of limited resolution and no side chains were modeled for this region.

### KCC3 architecture and dimerization

The KCC3 structure is captured in an inward-facing conformation, sharing the same LeuT-fold as other APC superfamily members (Figure 1A-C). TM1-5 and TM6-10 are organized in pseudo-symmetry, with TM3-5 and TM8-10 forming the scaffold domain and TM1, 2, 6, 7 constituting the bundle domain. A N-terminal loop is located between the scaffold and buddle domain (Figure 1D). TM domain and CTD are connected through the Scissor Helix likewise in *Dr*NKCC1 (39). The connection is stabilized by a salt bridge between the Scissor Helix and TM12 extension that are linked via a shorter hinge region (Supplemental Figure. S3A), resulting in a more rigid structure than *Dr*NKCC1. The PHSs are adhered to the peripheral region of CTD (Figure 1D). A huge displacement is observed in KCC3 CTDs compared with *Dr*NKCC1, which rotates about 70° clockwise as seen from cytosolic side when TM domains aligned together (Supplemental Figure S3B), revealing differences of domain arrangement among CCCs. The ECD of the two protomers are placed in distance, consisting of a long loop between TM5-TM6 and a short loop between TM7-TM8. The CCCs folds similarly around extracellular gate (Supplemental Figure S3C), interacts with TM4 and TM10 by salt bridges (Supplemental Figure S3D). This suggests a shared ECD gating mechanism in LeuT-fold transporters (40–42).

The two protomers assemble in domain-swapping conformation just like *Dr*NKCC1 (43–45). TM domain, Scissor Helixes and CTD are involved in dimerization of the transporter. Different from KCC1, the ECD of KCC3 are placed in distance (Figure 1B). The TM domains contact each other through several aromatic residues in TM11 and TM12. TM12 is spaced apart by the Scissor Helixes at the cytosolic side, tilting of which results in a large cleft between TM domains, which will be referred as Central Cleft hereafter (Figure 1D). The Central Cleft is surrounded by TM10-12, isolated from cytoplasmic environment by Scissor Helix (Supplemental Figure S3E). Unlike *Dr*NKCC1 or KCC1, no stable bound lipids or detergents are observed in the Central Cleft. Both the hydrophilic interactions between the Scissor Helixes (Supplemental Figure S3F) and the hydrophobic interactions between CTD and Scissor Helix from another protomer (Supplemental Figure S3G) contribute to dimerization.

### N-terminal loop in the cytosolic entry inhibits transport activity

A unique N-terminal loop encompassing N100-N120 is found to be bound in the entry of the cytosolic vestibule in the cryo-EM map (Figure 2A). The corresponding N-terminal loop of KCC2 encompassing A65-N83 is also found to be bound in the same location with the same mode, consisting with the sequence conservation in this region (Figure 2B, Supplemental Figure S4A). Focused 3D classification revealed a class in which this N-terminal loop is almost absent (Supplemental Figure S4B). This loop can be divided into N and C lobes, lying in the funnel formed by TM1, TM5, and TM8, between the bundle and scaffold domain. The N lobe is anchored between the TM1a, TM5, and IL23 loop through electrostatic interaction (Figure 2A). Following the bending at L108, the C lobe extends along TM8 and approaches CTD, stabilized through salt bridges E116-K549 and E119-R886, as well as hydrophilic interaction between H115 and F203 from IL23 (Figure 2A).

**Figure 2.**
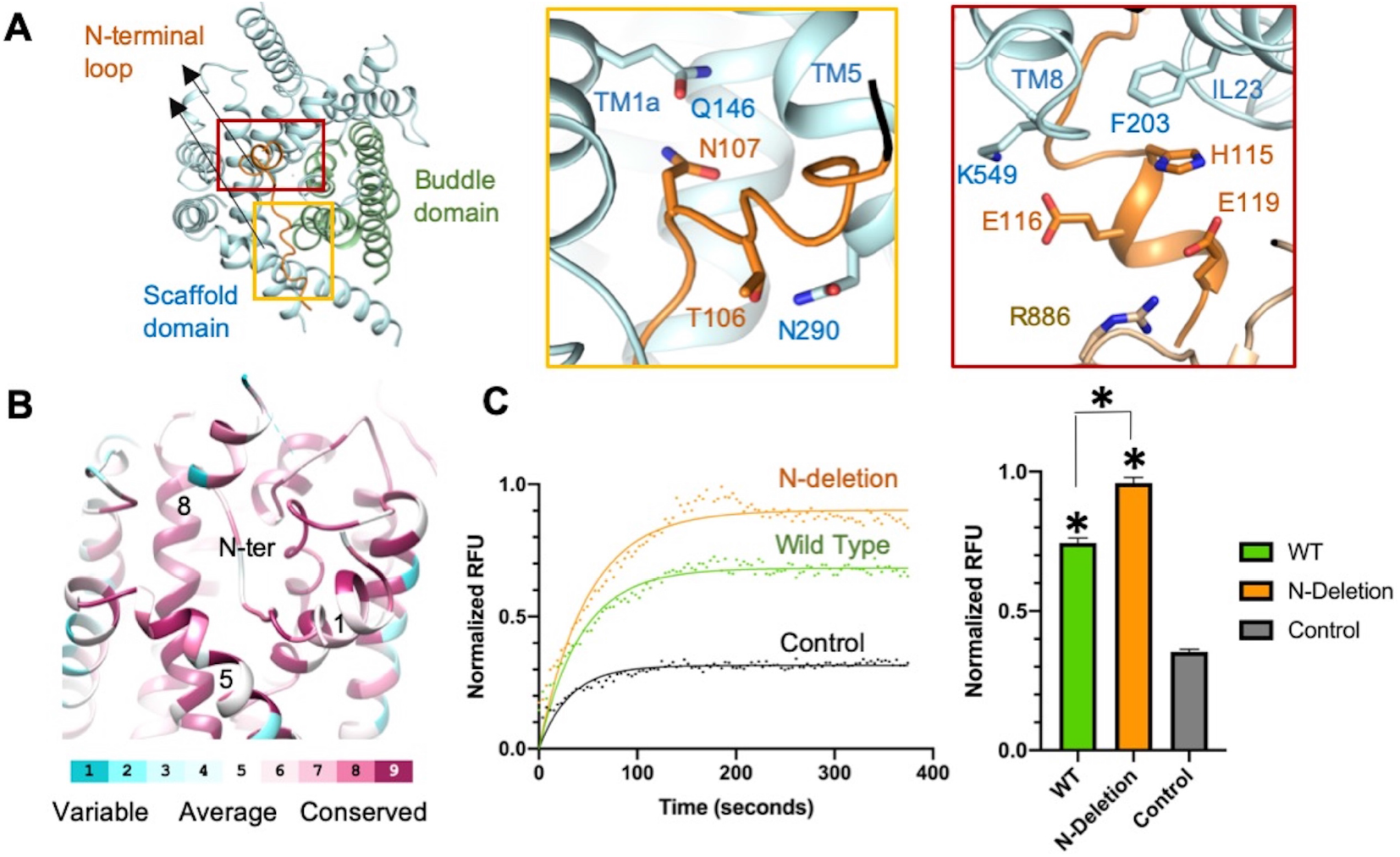
The inhibitory N-terminal loop of KCC3. (A) N-terminal loop locates in the cytosolic entry of the transport path of KCC3, between Buddle domain and Scaffold domain. Yellow inset: N lobe of N-terminal loop binding through interaction with TM1a, and TM5. Scarlet inset: C lobe bound to TM8 extension and CTD. (B) The conservation results around N-terminal loop calculated by Consurf (64, 65). The high conservation regions are labeled in red, while region with low conservation are labeled in cyan. (C) N-deletion has higher transport activity than Wild Type. The fluorescence readout as a function of time of different groups (Orange for N-deletion, Green for Wild Type, Grey for control, n=3 for every group) are plotted and fit into one-phase exponential curve. The average of last 23 measurements are plotted. Statistic differences are estimated by one-way Anova (*p<0.0001).

The N-terminal loop is the inhibitory regulation element of KCCs. The binding locks the transporter in inward-facing state, prohibits the conformation change of bundle domain. At the same time, it seals up the cytosolic entry (Supplemental Figure S4C). The loop binding also leads to more electronegativity in the entry, reducing its attraction to the negative-charged chloride ions (Supplemental Figure S4D). The high sequence conservation of the binding pocket for N-terminal loop suggests the functional significance (Figure 2B). To investigate its role, a KCC3 mutant with deletion of the N-terminal loop (N-Deletion) is designed. Compared with wild type KCC3 (46), N-Deletion mutant exhibits higher transport activity (Figure 2C), supporting the auto-inhibitory effect of the N-terminal loop.

### Phosphorylation harboring segments in CTD

Three conserved residues S45, T940 and T997 in KCC3 can be phosphorylated by the same kinase and are well studied (26). The S45 residue is located in the most N-terminal loop that is invisible in the cryo-EM map. The segments 931-972 and 994-1000 of KCC3 containing T940 and T997 were resolved in the cryo-EM map (Figure 3A). For simplicity, these two segments are named as phosphorylation-harboring segment 1 and 2 (PHS1 and PHS2), respectively. 3D classification focused on PHSs region revealed two classes. The density for PHSs is well resolved in class 1, while almost invisible in class 2 (Supplemental Figure S5A). In order to investigate the role of phosphorylation, we generated two mutants, one of which imitates the phosphorylation status with mutation S45E/T940E/T997E, while the other mimics dephosphorylation status with mutation S45A/T940V/T997V. These two mutants were designated as P-mimic and DP-mimic respectively for simplicity. We purified the mutant protein and solved the structures at resolution of up to 2.9 Å and 3.2 Å for P-mimic and DP-mimic mutant, respectively (Supplemental Figure S1, 2). 3D classification focused on PHSs region in both P-mimic and DP-mimic mutants revealed two classes similar to that for the WT KCC3. The solved PHS1 density in P-mimic is similar to that in WT KCC3.

**Figure 3.**
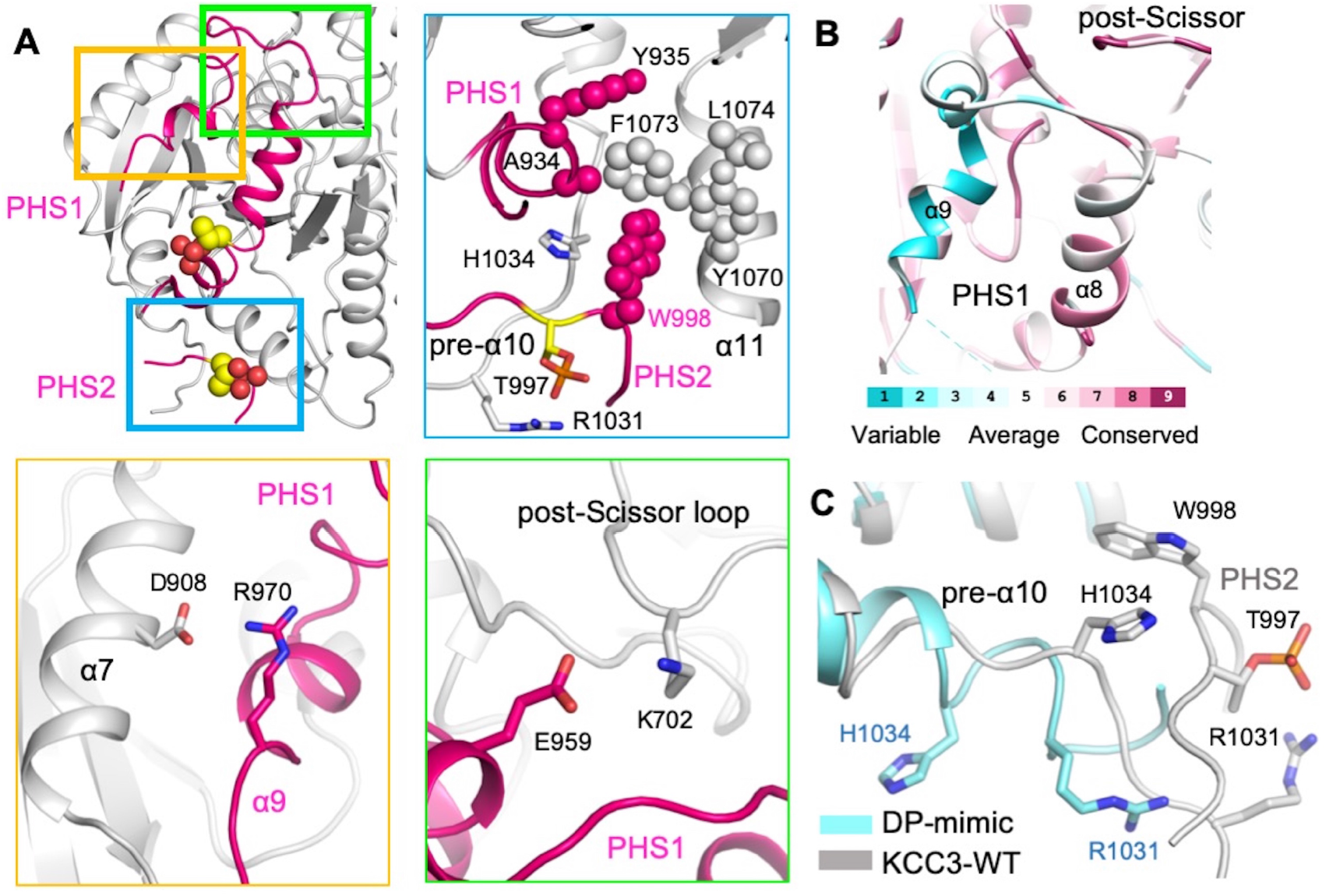
Phosphorylation sites in KCC3. (A) Overview of the phosphorylation sites in KCC3 structure. Both of the visible phosphorylation sites are located in the peripheral area of the CTD. The main interaction regions are highlighted by rectangles. Blue subset: Both hydrophobic interaction and hydrophilic interaction participates in PHS2 binding. PHS2 binds to a hydrophobic pocket through W998. Yellow subset: Salt bridges between α7 of CTD and α9 of PHS1. Green subset: Salt bridges between post-Scissor loop and α8-9 loop of PHS1. (B) Consurf results of PHS1 region. The conservation of residues is labeled as indicated. (C) Structure rearrangement of pre-α10 loop. The resulted different orientation of the H1034 and R1031 ceases the binding of PHS2.

However, the solved density for PHS1 in DP-mimic is less order than that in WT, suggesting that the solved PHS1 segment of WT protein is in phosphorylated status (Supplemental Figure S5A), which is further supported by Mass Spectra results (Supplemental Figure S5B). The status of PHSs is affected by the phosphorylation of T997. The phosphorylated T997 facilitates the binding of PHS2 by interacting with R1031 in the pre-α10 loop. A conserved W998 in PHS2 is inserted into a hydrophobic pocket constituted from N-terminal end of PHS1, α11 of CTD, and pre-α10 loop and further stabilized by H1034 from pre-α10 loop via π-π interaction (Figure 3A). The hydrophobic pocket helps to stabilize PHS1. The PHS1 further interacts with CTD through salt bridge between E959 on PHS1 and K702 from post-Scissor loop, R970 on PHS1 and D908 from α7 (Figure 3A). As these two salt bridges locate near α9 in PHS1, the lack of sequence conservation around the region marks the particular function in KCCs (Figure 3B), which is partially supported by the different phosphorylation effects of CCCs (11). PHS1 adopts a significant different position than its counterpart in *Dr*NKCC1 (Supplemental Figure S3B), suggesting different regulation mechanism between KCCs and NKCCs. In DP-mimic, a significant conformation change is observed in pre-α10 loop that is rearranged as a helix with H1034 pointing away from hydrophobic core, disrupting the interaction with PHS2 (Figure 3C). This hampers PHS1 adhesion to CTD, coordinates with the 3D classification results (Supplemental Figure S5A).

### DIOA inhibits transport activity by anchoring TM10

[(dihydroindenyl)oxy] alkanoic acid (DIOA) was firstly identified as a specific inhibitor targeting NEM-stimulated and bumetanide-insensitive K^+^ efflux in human red cells (36). DIOA is widely applied in KCCs research and has potential application in drug development. In order to investigate the inhibition mechanism of DIOA to KCCs, the structure of KCC3 in complex with DIOA was solved at resolution up to 2.5 Å, in which two non-protein densities are observed in the Central Cleft and each built as DIOA as the density shape is well fit with DIOA (Figure 4A).

**Figure 4.**
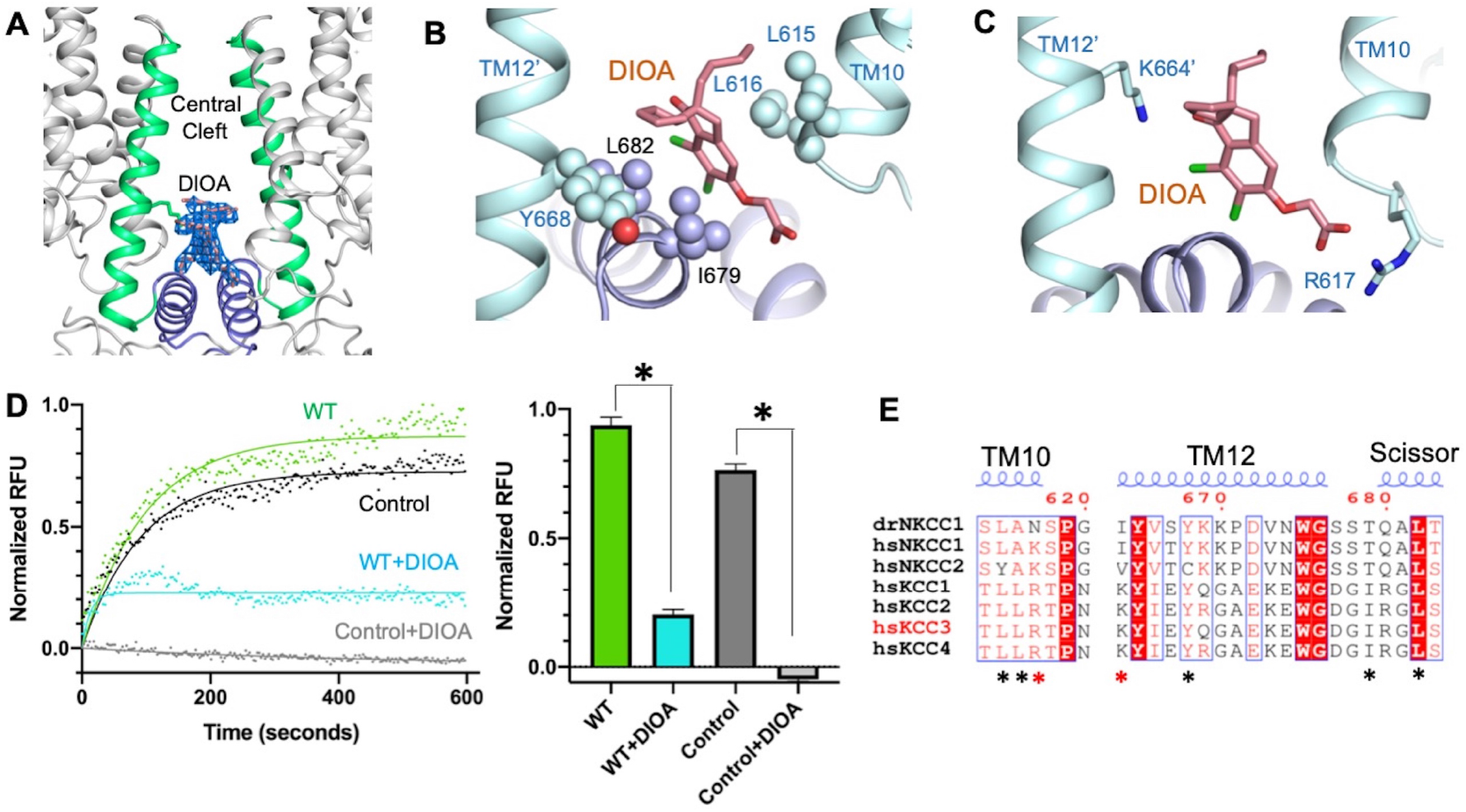
DIOA binding details in KCC3. (A) DIOA locates in the Central Cleft. The density is well-fit for DIOA. (B) Hydrophobic interaction from TM10, Scissor Helix and TM12’ stabilizes DIOA. (C) Hydrophilic interaction from TM10 and TM12’. The interaction with R617 stabilizes the conformation of TM10. (D) DIOA inhibits the transport activity of KCC3 expression cells. The fluorescence readout as a function of time of different groups (Green for Wild Type, Cyan for Wild Type with DIOA incubation, Dark grey for control, Light grey for control incubated with DIOA. n=4 for every group) are plotted and fit into one-phase exponential curve. The average of last 30 measurements are plotted. Statistic differences are estimated by one-way Anova (*p<0.0001). (E) Multi-alignment of the key residues involved in DIOA binding. Although R617 mutate to K in NKCCs retains the charge of sidechain, K664 mutate to I/V mostly stops the binding with DIOA. Along with the absence of Central Cleft, NKCCs therefore are insensitive to DIOA.

DIOA is sandwiched by the hydrophobic residues L682 and I679 from Scissor Helix, as well as L612, L615, L616 from TM10 (Figure 4B). DIOA also interacts with TM12 and TM10-11 loop via hydrophilic interactions with K664 in TM12 and R617 in TM10-11 loop (Figure 4C). These interactions may prohibit the potential movement of TM10 in the transport cycle of the transporter and lock KCC3 in the inward-facing conformation. The inhibitory effect is observed in various research and cell-based assay (Figure 4D). The key residues involved in DIOA binding are mostly conserved in KCCs but not in NKCCs (Figure 4E). Moreover, the Center Cleft is absent in *Dr*NKCC1 or KCC1. All of these suggest the specificity of DIOA to KCC2 and KCC3.

## Discussion

In this work, we solved the structures of the full-length human KCC3 and KCC2, and investigated multiple regulatory elements from protein *per se* or from specific chemicals, provided more information on the regulation of KCCs (Figure 5A).

**Figure 5.**
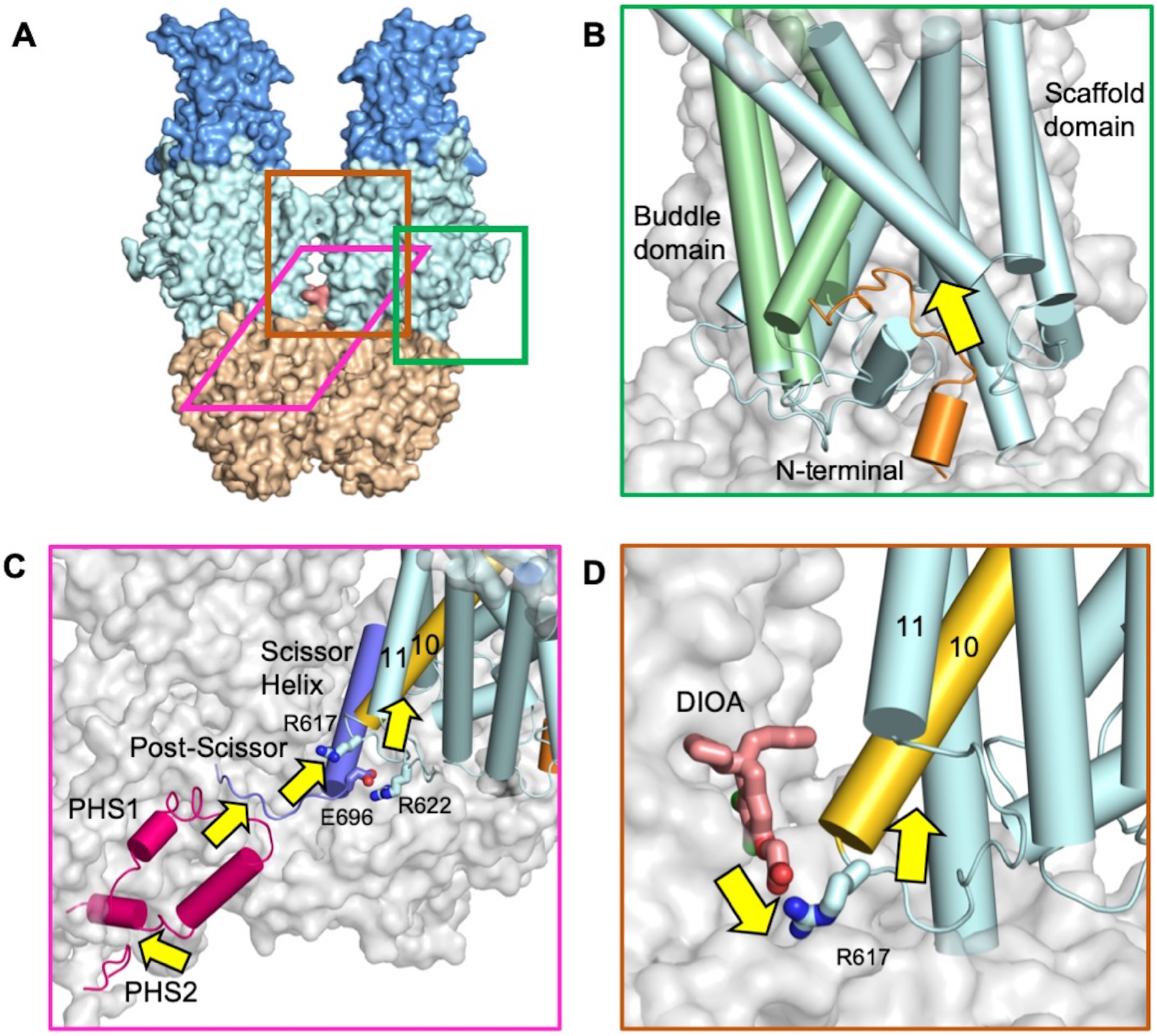
Diagram scheme for the regulation model of the discussed elements. (A) The region involved in different regulatory elements are plotted in the structure of KCC3. The structure model is colored by domains. (B) N-terminal inhibition of KCC3 activity. N-terminal blocks the cytosolic vestibule of the co-transporter. Both space occupation and charge alternation by N-terminal loop prohibits ion transport. (C) Phosphorylation facilitated PHSs binding may stabilize inward-facing conformation of KCC3 through interaction with post-Scissor loop. As E696 in post-Scissor loop is located closely with two Args from TM10-11 loop, the possible interaction in between may help in stabilization of TM10 movement and further lock the co-transporter in inward-facing conformation. (D) DIOA locks TM10 conformation by directly interaction with R617 of TM10-11. The locking of TM10 stabilizes KCC3 in inward-facing conformation and therefore inhibits the transport activity.

The N-terminal loop is found to be bound in the cytosolic entry of KCC3, which block the transport path and prohibit the possible conformation change during transport cycle (Figure 5B). The inhibitory effect is proved by our transport assay that showed the deletion of the N-terminal loop increased the transport activity. The blockage role of N-terminal loop may avoid the unexpected potential ion leakage through KCCs *in vivo*. The N-terminal deletion in KCC2 was identified in Epilepsy of infancy with migrating focal seizures patients, with the deletion alone was sufficient to induce Cl^−^ transport abnormality and no influence on protein location (47), highlighting the importance of N-terminal loop for transport activity. The inhibitory binding of the N-terminal loop represents an example for the regulation of the transporter activity mediated by peptides, providing structural basis for the future development of the peptide drugs (48, 49). Interestingly, the N-terminal sequence is partially retained in NKCC2, but absent in NKCC1 (Figure 2B), which may facilitate development of the homolog-specific peptide-drug.

Phosphorylation is one of the most important post-translation modifications of the CCCs. There are multiple kinases participating in the process (27, 50, 51) that target to different residues. In this work, we focus on the PHS region that contains phosphorylated sites targeted by WNKs pathway. Here we proposed a model for the regulation mechanism of phosphorylation (Figure 5C). The PHSs are stabilized through the hydrophobic core formed by N-terminal end of PHS1, W998 of PHS2 and α11 of CTD. The tip of PHS1 is located closely to and interacts with the post-Scissor loop through a salt bridge between its E959 and K702 of the post-Scissor loop, which pulls the TM10-11 loop through E696 that may interact with R622 and/or the main atoms on the TM10-11 loop. The phosphorylation of T997 enhances its interaction with R1031 on pre-α10 loop and may stabilize PHSs binding, which might inhibit the transport activity by stabilizing the inward-facing conformation through stabilizing TM10-11 loop via interactions between the tip of PHS1 and post-Scissor loop (Figure 5C), providing a possible answer for the phosphorylation-inhibition mechanism of KCCs. Mutation in the corresponding residue in KCC2 of R1031 of KCC3 results in epilepsy(52), suggesting its important role.

In DP-mimic mutant, dephosphorylation accompanied with the conformation change of the pre-α10 loop impedes the formation of the hydrophobic core and the binding of PHSs, leading to less-visible and less-ordered PHSs density in the cryo EM map. The post-Scissor loop contains multiple conserved prolines, suggesting the loop structure of this region is critical for the regulation mechanism discussed here. This long-range allosteric regulation from PHSs requires involvement of multiple elements. All of these may explain the heterogeneity in this region revealed by 3D classification. Multiple disease-related truncations are identified around the CTD and PHSs, suggesting the important role of these regions. This inhibitory mechanism is similar to that for DIOA, in which the locking or stabilizing of the TM10-11 loop is crucial for the inhibitory effect (Figure 5D).

## Methods

### Expression and purification of KCC2, KCC3, KCC3 phosphorylation site mutants, and KCC3 N-deletion

The protein expression and purification process of the proteins are alike. The human full length KCC2b was purchased from Jingmai Biotech Inc. KCC3b sequence were amplified from cDNA library. DP-mimic and P-mimic were introduced to the KCC3 sequence by a standard two-step PCR. KCC3 N-deletion was designed based on the structural information, with 101-120 amino acids deleted, and accomplished by standard two-step PCR. All protein encoding sequences were subcloned to pCAG vector with N-terminal FLAG-tag for expression in HEK293F cell lines transfected by polyethylenimines (PEIs, Polysciences). Approximately 1.5 mg plasmids were incubated with 3 mg PEIs in 50 mL SMM 293T-I medium (Sino Biological Inc.) for 30 min before transfection. Cells were cultured in Multitron-Pro shaker (Infors, 130 rpm) under 37° for 48hs before collection and resuspension in Lysis Buffer (25 mM Tris 8.0 and 150 mM NaCl). Protease inhibitors containing 1.3 mg/ml aprotinin, 5 mg/ml leupeptin, 1 mg/ml pepstanin (Amresco), and 0.2 mM PMSF (Sigma) was added before cruel-extraction at 4°C in 1.5% DDM and 0.3% CHS (Anatrace) for 2 hours. After centrifugation at ~18000g for 1 hour, the supernatant was applied to anti-FLAG M2 affinity gel (Sigma). Then washed by Wash buffer which is made up of Lysis buffer supplied with protease inhibitors and 0.02% GDN (Anatrace), the protein was eluted by 0.2mg/ml FLAG peptide. Finally, the protein was concentrated by 50-kDa cut-off Centricon (Millipore) and performed size exclusion chromatography (SEC, Superose 6, 10/300, GE Healthcare) in Wash buffer. The peak fractions were confirmed through SDS-PAGE analysis and concentrated for Cryo-EM sample preparation.

### Cell-based assay estimating N-terminal loop function in transport

FluxOR II Green Potassium Ion Channel Assay kit (Invitrogen) is applied to estimate the transport activity of KCC3 WT and N-deletion proteins. Cells of SF9 were transfected at 2 × 106 cell/mL by baculovirus encoding KCC3 WT and N-deletion. Control group was uninfected. After 48 hours suspension culturing, cells were placed in plate for 2 hours allowing adhesion. 1 × Loading buffer replaced medium and incubated with the cells in 27 °C for 1 hour. After treatment by low-osmotic buffer (88 mM NaCl, 42 mM NMDG-Cl, 5 mM KCl, 2 mM CaCl2, 1 mM MgCl2, 0.77 mg/ml Probenecid, 10 mM HEPES, pH 7.4, osmolarity 260 mOsm/kg) for 45 min, cells were incubated in Assay Buffer ready for fluorescence measurement by Varioskan LUX microplate reader (Thermo Fisher). For measuring the effect of DIOA on transport activity, 100uM DIOA was added during low-osmotic treatment. The excitation wave length was ~488 nm with width of 12 nm, and emission at 525 nm was recorded. The average of first 590s measurement act as the baseline. Data was recorded 10 seconds after addition of Basal Potassium Stimulus buffer, and recorded every 3s for about 375s, then every 10s for 230s.

### Cryo-EM sample preparation and data acquisition

To prepare cryo-EM samples, aliquots (3 μL) of the purified protein concentrated to ~ 10 mg/mL were placed on glow-discharged holey carbon grids (Quantifoil Au R1.2/1.3). For KCC3-DIOA samples, 500 μM DIOA was incubated with the protein solution for 2 hs before concentrated for cryo-EM sample preparation. The grids were blotted for 5 s and flash-frozen in liquid ethane cooled by liquid nitrogen with Vitrobot (Mark IV, Thermo Fisher Scientific). The prepared grids were transferred to a Titan Krios operating at 300 kV equipped with Gatan K3 detector and GIF Quantum energy filter. Movie stacks were automatically collected using AutoEMation (53) with a slit width of 20 eV on the energy filter and a defocus range from −2.2 μm to −1.2 μm in super-resolution mode at a nominal magnification of 81,000×. Each stack was exposed for 2.56 s with an exposure time of 0.08 s per frame, resulting in a total of 32 frames per stack. The total dose rate was approximately 50 e^−^/Å^2^ for each stack. The stacks were motion corrected with MotionCor2 (54) and binned 2-fold, resulting in a pixel size of 1.087 Å/pixel. Meanwhile, dose weighting (55) was performed. The defocus values were estimated with Gctf (56).

### Data processing

Cryo-EM data was processed similarly for all protein samples. Particles were automatically picked using Relion 3 (57) from manually selected micrographs. After 2D classification, good particles were selected and subjected to global angular searching 3D classification against an initial model generated with Relion 3 with C2 symmetry. For each of the last several iterations of the global angular searching 3D classification, a local angular searching 3D classification was performed, during which the particles were classified into 4 classes. Non-redundant good particles were selected from the local angular searching 3D classification. Then, these selected particles were subjected to multi-reference 3D classification, local defocus correction (56), 3D auto-refinement and post-processing. To further improve the map quality, the dataset was C2 symmetry-expanded and then classified with adapted masks applied on the intracellular domain (CTD) and the TM/ECD, respectively. The particles with N-terminal loop bound or the particles with well-ordered PHS were selected and further focused refined with appropriate mask.

The 2D classification, 3D classification and auto-refinement were performed with Relion 3. The resolution was estimated with the gold-standard Fourier shell correlation 0.143 criterion (58, 59) with high-resolution noise substitution (60). Refer to Extended data fig. 1-2 and Extended Data Table 1 for details of the data collection and processing.

### Model building and structure refinement

Model building of KCC3 was performed *ab initio* with Phenix (61) and Coot (46) based on the focused-refined cryo-EM maps with aromatic residues as land markers, as most of these residues were clearly visible in our cryo-EM map. Each residue was manually checked with the chemical properties considered during model building. Several segments of the sequence were not modeled because of the invisibility of the corresponding density in the map. The model building of P-mimic, DP-mimic and KCC2 was accomplished with similar methods using the KCC3 model as a starting template. Chainsaw in CCP4 (62) package was used to substitute the sequence during the model building of KCC2.

Structure refinement was performed with Phenix (61) with secondary structure and geometry restraints to prevent structure overfitting. To monitor the overfitting of the model, the model was refined against one of the two independent half maps from the gold-standard 3D refinement approach. Then, the refined model was tested against the other map (63). Statistics associated with data collection, 3D reconstruction and model building can be found in Supplemental Table S1.

## Acknowledgement

We thank the cryo-EM facility, the supercomputer center and the mass spectrometry & metabolomics core facility of Westlake University for providing cryo EM, computing and mass spectrometry support, respectively. This work was supported by the National Natural Science Foundation of China (projects 31971123, 81920108015, 31930059), the Key R&D Program of Zhejiang Province (2020C04001).

