## Supplemental for "Molecular basis for regulation of human potassium chloride cotransporters"

**Table S1 | Data collection, 3D reconstruction and model statistic**

| | | | | | |
|---|---|---|---|---|---|
| **Data collection** | | | | | |
| EM equipment | Titan Krios (Thermo Fisher Scientific) | | | | |
| Voltage (kV) | 300 | | | | |
| Detector | Gatan K3 Summit | | | | |
| Energy filter | Gatan GIF Quantum, 20 eV slit | | | | |
| Pixel size (Å) | 1.087 | | | | |
| Electron dose (e-/Å2) | 50 | | | | |
| Defocus range (μm) | -1.2 ~ -2.2 | | | | |
| Sample | KCC3 | KCC2 | KCC3 (P-mimic) | KCC3 (DP-mimic) | KCC3-DIOA |
| Number of collected micrographs | 4,488 | 2,184 | 3,914 | 4,589 | 3,460 |
| Number of selected micrographs | 3,783 | 1,862 | 2,511 | 1,988 | 3,201 |
| **3D Reconstruction** | | | | | |
| Software | Relion 3.0 | | | | |
| Number of used particles (Overall) | 453,155 | 186,236 | 345,037 | 115,055 | 364,959 |
| Resolution (Å) | | | | | |
| Overall | 3.3 | 3.2 | 3.1 | 3.6 | 2.7 |
| TM_Extracellular | 3.1 | 2.9 | 3 | 3.3 | 2.7 |
| Intracellular | 2.9 | 2.9 | 2.9 | 3.2 | 2.8 |
| Symmetry | Overall: C2; TM_Extracellular: C1; Intracellular: C2 | | | | |
| Map sharpening B-factor (Å2) | Overall: -90; TM_Extracellular: -150; Intracellular: -90 | | | | |
| **Refinement** | | | | | |
| Software | Phenix | | | | |
| Cell dimensions | | | | | |
| a=b=c (Å) | 347.84 | 347.84 | 347.84 | 347.84 | 347.84 |
| α=β=γ (°) | 90 | 90 | 90 | 90 | 90 |
| Model composition | | | | | |
| Protein residues | 1,876 | 1,872 | 1,868 | 1,866 | 1770 |
| Side chains assigned | 1,876 | 1,850 | 1,868 | 1,798 | 1770 |
| Sugar | 16 | 14 | 16 | 16 | 16 |
| ligand | 2 | 2 | 2 | 2 | 2 |
| K+ | 2 | 2 | 2 | 2 | 2 |
| Cl- | 4 | 2 | 4 | 4 | 4 |
| inhibitor | | | | | 2 |
| Water | 2 | 2 | 22 | 2 | 40 |
| R.m.s deviations | | | | | |
| Bonds length (Å) | 0.005 | 0.009 | 0.007 | 0.009 | 0.006 |
| Bonds Angle (°) | 0.967 | 1.155 | 0.938 | 1.068 | 1.006 |
| Ramachandran plot statistics (%) | | | | | |
| Preferred | 89.29 | 88.19 | 91.68 | 87.65 | 91.75 |
| Allowed | 10.28 | 10.94 | 8.1 | 12.03 | 7.91 |
| Outlier | 0.43 | 0.87 | 0.22 | 0.32 | 0.34 |

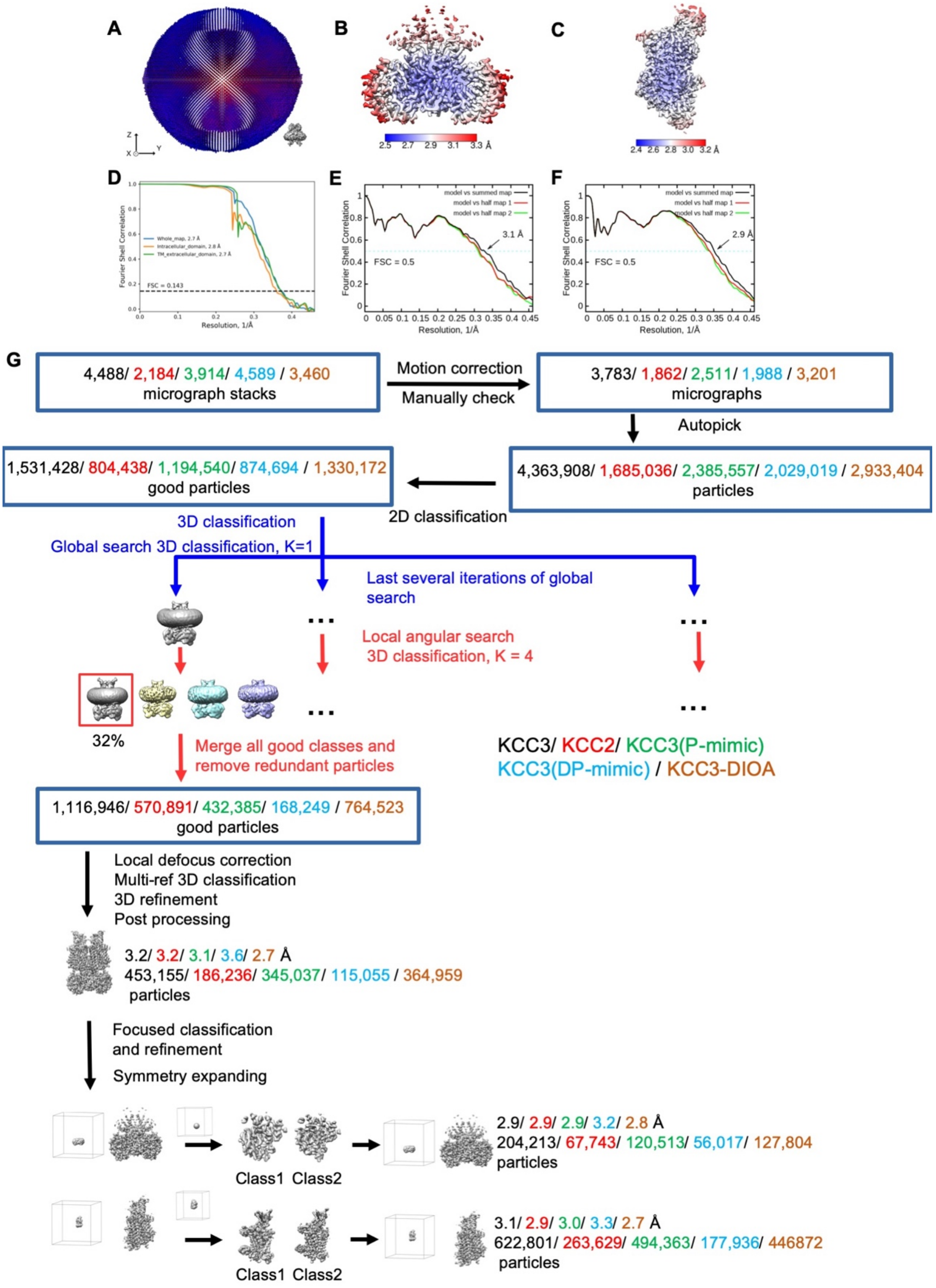


**Figure S1. Cryo-EM analysis of KCC3, KCC2, KCC3(P-mimic), KCC3(DP-mimic) and KCC3-DIOA.**

(A) Euler angle distribution of KCC3.

(B and C) Local resolution maps for the 3D reconstruction of KCC3 focused on the intracellular domain or TM region and extracellular domain, respectively.

(D) Gold standard FSC curve for the 3D refinement of the overall structure, and the 3D refinements focused on the intracellular domain and TM region and extracellular domain, respectively.

(E) FSC curve of the refined model of KCC3 versus the intracellular domain map that it is refined against; of the model refined against the first half map versus the same map (red); and of the model refined against the first half map versus the second half map (green). The small difference between the red and green curves indicates that the refinement of the atomic coordinates did not suffer from overfitting.

(F) is same to (E), but for the TM/ECD region of KCC3.

(G) Flowchart for cryo-EM data processing. Please refer to the ‘Data Processing’ in Methods section for more details.

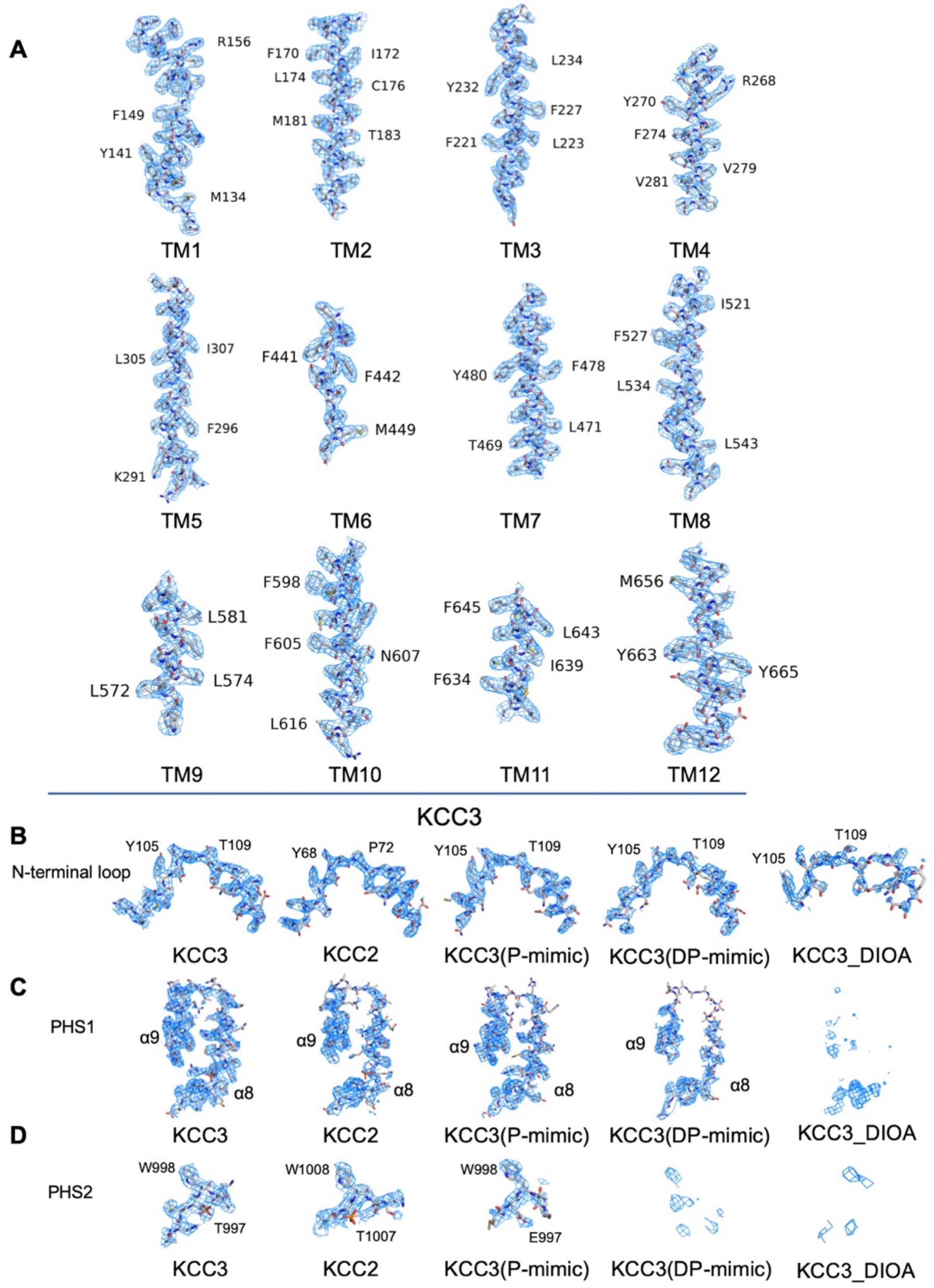


**Figure S2. Cryo EM density maps.**

(A) Cryo EM density maps for TM segments of KCC3 shown at threshold of 8 σ.

(B) Cryo EM density maps of N-terminal loop shown at threshold of 12 σ.

(C) Cryo EM density maps of PHS1 region shown at threshold of 8 σ.

(D) Cryo EM density maps of PHS2 region shown at threshold of 8 σ.

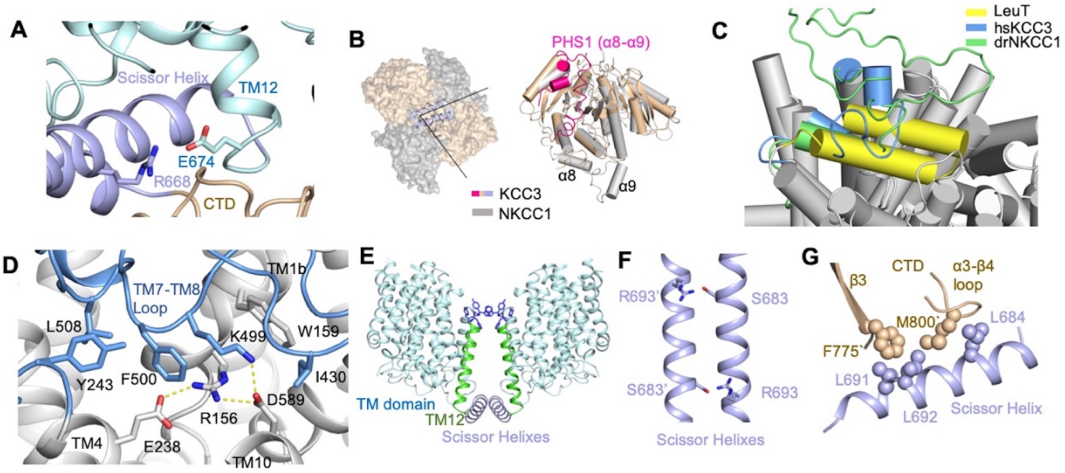


**Figure S3. Domain arrangement of KCC3.**

(A) Interaction between Scissor helix and TM12 extension. The limited loop region between TM and scissor helix corelates the little flexibility of CTD conformation in KCC3.

(B) Structure compare of drNKCC1 (PDB ID: 6npl) and KCC3. Left: superpose CTD domain of drNKCC1 and KCC3. Approximately 70° rotation are observed. The angle is placed along Scissor Helix. Right: superpose CTD from one protomer, highlighting the different conformation of PHS1 in the two structures.

(C) The high similarity around the ECD of KCC3 (this study), drNKCC (PDB ID: 6npl), and LeuT (PDB ID: 3TT3).

(D) The salt bridges (E238-R156-D589-D499) around the extracellular region of the KCC3 act as a conserved gate.

(E) Aromatic residues conduct the dimer interface between TM domains. TM12 in KCC3 are pushed away by scissor helixes insertion.

(F) Hydrophilic interactions between R693 and S683’ in Scissor Helix participate in dimerization. Single prime presents the residues from another protomer.

(G) Interaction between Scissor helix and CTD. Hydrophobic residues participate in the interactions are represented in sphere.

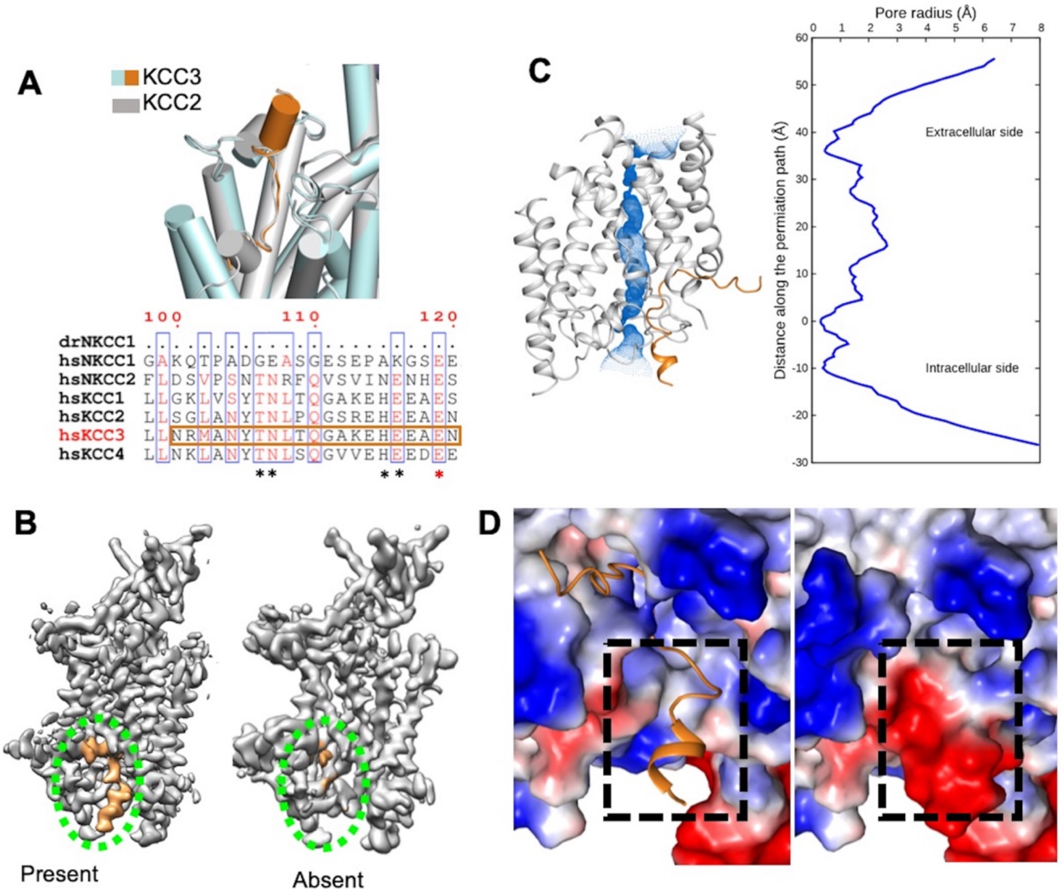


**Figure S4. N-terminal loop binding in KCC3.**

(A) The N-terminal loop structure is conserved in KCC2 and KCC3. The key binding residues are conserved in KCCs, and highlighted by stars. Red star presents higher conservation level.

(B) Two classes of N-terminal loop region can be detected in the 3D classification. Electron density of N-terminal loop is highlighted in orange.

(C) N-terminal loop binding narrows down the inward-open entry. Permeation path calculated by HOLE (1). Left: blue dots represent the permeation path of KCC3. Right: The pore radii along the conducting passage.

(D) Surface electrostatic potential of TM domain calculated by PYMOL (2). Dashed line rectangle highlights the N-terminal loop binding region to show the alternation of electrostatic potential.

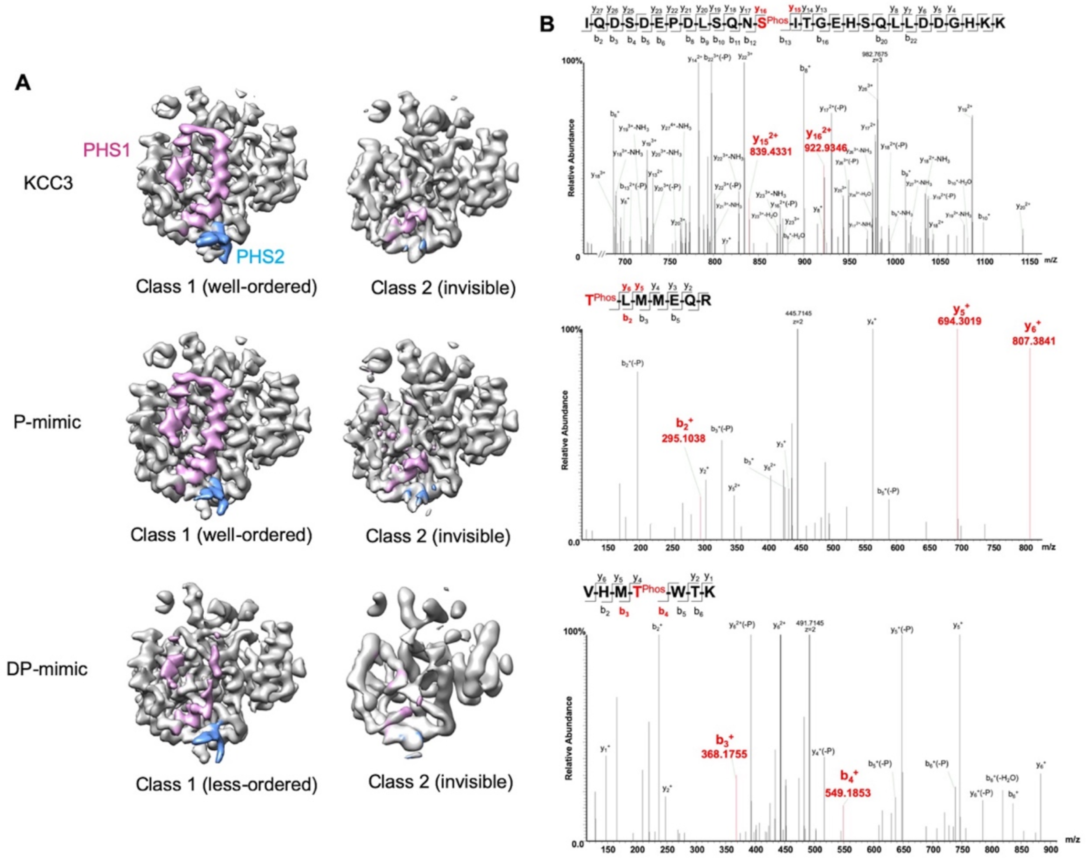


**Figure S5. Density of phosphorylation harboring sites.**

(A) Cryo-EM density map focused on CTD of KCC3, P-mimic, and DP-mimic are represented. Density of PHS1 and PHS2 are labeled by different colors. While well-ordered classes are presented in both wild type and P-mimic, a class of less-ordered PHS1 was observed in DP-mimic.

(B) Mass Spectra result of KCC3 WT protein phosphorylation status. Top: S45 is phosphorylated as shown by molecular weight (MW) difference between $y_{15}^{2+}$ and $y_{16}^{2+}$. Middle, T940 is in the top of the charged peptide. Its phosphorylated state is illustrated by both MW difference between $y_5^{2+}$ and $y_6^{2+}$, as well as dephosphorylated state of L941 by $b_2^{+}$. Bottom: T997 is phosphorylated as supported by MW difference of $b_3^{+}$ and $b_4^{+}$.
